# Ephaptic coupling improves the neural population code

**DOI:** 10.64898/2026.07.28.741366

**Authors:** Kianoush Banaie Boroujeni, Sabine Kastner

## Abstract

Neurons share an extracellular medium, so every spike’s current sets up a local field that feeds back onto the membrane potential of nearby cells, an interaction known as ephaptic coupling. The local field is usually studied by imposing it externally, for example by injecting sinusoidal currents through an extracellular electrode, which entrains and synchronizes firing. However, an endogenous field, self-generated by the network’s own spiking, is a different phenomenon whose functional role stays unclear, because it cannot be removed without silencing the neurons that produce it. We built matched spiking networks with and without a self-consistent endogenous field, and a rate-matched control separating the field from firing rate. The results show that the origin of the field determines the effect. A field the network generated for itself decorrelated the population, expanded its dimensionality, and improved decoding, whereas the same field imposed from outside did the opposite, synchronizing the population and reducing coding performance. The benefit held across a spatial and a temporal coding task and survived rate-matching. A closed-form theory connects the decorrelation to the field acting as a spatial filter on the slow component that neurons share. To look for this in cortex, we analyzed multi-patch recordings, showing that a neuron’s spikes left a distance-dependent trace in unconnected neighbors which we reproduced by the same field model. The self-generated field is therefore not a by-product but a mechanism that, without wiring, can improve the population code and reshape the geometry of neural population dynamics.

## Introduction

Ephaptic coupling is the process by which the electric field generated by neural activity acts back on the membranes of neighboring neurons, without a chemical or electrical synapse (Arvanitaki, 1942; Jefferys, 1995). Because neurons share a resistive extracellular medium, the transmembrane currents of an active population generate a field that reaches every cell and shifts its potential, the same currents we record as the local field potential (Buzsaki et al., 2012). This field is usually treated as a by-product of activity, yet it feeds back onto the membranes that create it, so a network is always immersed in a field of its own making. The effect was demonstrated long ago, with the extracellular current of one spike shifting the excitability of an inactive neighbor and weak fields modulating an already firing cell far more readily than they excite a silent one (Katz and Schmitt, 1940; Terzuolo and Bullock, 1956).

Ephaptic coupling has been reported at different levels, from the synapse to brain-wide networks (Jefferys, 1995; Anastassiou and Koch, 2015). At the synaptic level, retinal horizontal cells feed back onto cone photoreceptors through the field they generate in the synaptic cleft rather than through a transmitter (Kamermans et al., 2001). At the single-neuron level, the field from one cell has been reported to directly inhibit a neighbor (Furukawa and Furshpan, 1963; Faber and Korn, 1989; Blot and Barbour, 2014; Han et al., 2020). At the circuit level, extracellular fields in cortex and hippocampus have been shown to shift the membrane potential, bias spike timing, and entrain firing across a population (Radman et al., 2007; Bikson et al., 2004; Francis et al., 2003; Deans et al., 2007; Anastassiou et al., 2010, 2011; Lee et al., 2024). At the network level, a self-generated field can propagate slow or epileptiform waves across a cut that blocks synapses and gap junctions (Zhang et al., 2014; Qiu et al., 2015; Chiang et al., 2019).

Whether this coupling serves a function in the cortex and in neural computation, beyond such special cases, remains unresolved (Anastassiou and Koch, 2015; Weiss and Faber, 2010). One of the biggest obstacles is methodological. Because the endogenous field is produced by the network’s own activity, it cannot be scaled or removed without changing the spiking that creates it, which limits any causal test of its function, and the field is weak enough that any effect can be attributed to the synaptic activity it accompanies. The main approach so far has been to impose a field from outside and measure how firing changes, which has been shown to entrain neurons and amplify an ongoing rhythm (Ozen et al., 2010; Krause et al., 2019; Liu et al., 2018; Reato et al., 2010). But an imposed field is exogenous even when it is delivered in closed loop from the recorded activity (Frohlich and McCormick, 2010), because it is applied from outside rather than arising in the tissue that generates it, so what the intrinsic closed-loop field does cannot be inferred from these stimulation studies.

Two open questions then follow. First, does a self-generated field, one that emerges from the population’s own spiking and feeds back onto the circuit that produces it, act like the same field imposed from outside, or differently? Second, does the field change what a population encodes, not only how fast or how synchronously it fires? What a downstream area can read out from a population is limited by its shared, correlated variability (Averbeck et al., 2006; Cohen and Kohn, 2011; Moreno-Bote et al., 2014), and a shared field, generated from and feeding back on the population’s own dynamics, is a natural way to shape exactly that variability (Stacey et al., 2015). Previous modeling has examined how the extracellular potential of one cell perturbs its neighbors (Holt and Koch, 1999; Goldwyn and Rinzel, 2016), and recent recordings suggest the cortical field may itself encode stimulus and memory content (Pinotsis and Miller, 2022, 2023), yet no study has examined what the self-generated field itself does to a population code.

Here we ask what a self-generated field does to population coding, and whether the origin of the field matters for this coding. To address this, we simulate a spiking network with and without ephaptic coupling, in which the field is generated self-consistently by the population and fed back onto every neuron. We then contrast this self-generated field with the same field imposed from an external source, holding firing rate constant so that any difference reflects the respective source of activity rather than excitability. We find that the origin of the field is a key factor. A field that the network generates for itself decorrelates the population and improves its code, whereas the same field imposed from outside synchronizes it and reduces the coding performance toward chance; the benefit reflects a reorganization of shared variability and is not solely explained by the accompanying change in firing rate.

## Results

### A network model of ephaptic coupling

We placed 500 leaky integrate-and-fire (LIF) neurons, 400 excitatory and 100 inhibitory, at quasi-random positions on a two-dimensional sheet so that pairwise distances spanned the full extent of the array (**Figure 1a**). On top of the standard membrane equation, each neuron received an ephaptic term, a distance-weighted sum of its neighbors’ departures from rest scaled by a coupling strength that we express as a fraction of the value at which the feedback becomes critical (**Figure 1b**). This closed-loop model represents the ephaptic LIF (eLIF) network, in which setting the coupling to zero recovers the field-free LIF network. Each neuron in the network receives a baseline input, private noise, and a distance-dependent correlated noise input from external noise sources (**Figure S1a**).

**Figure 1.**
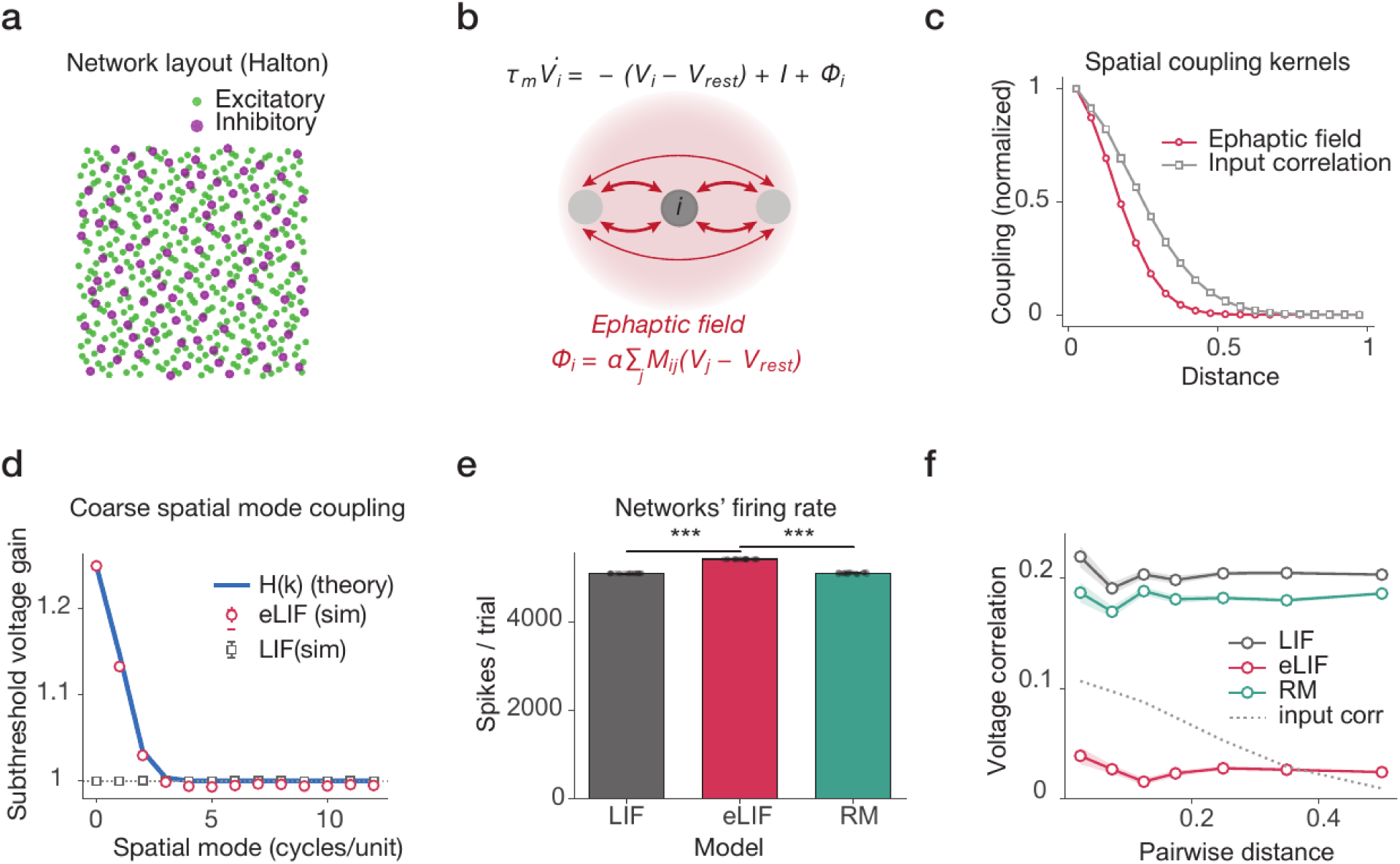
The eLIF network model and its effect on subthreshold voltage correlations. **(a)** positions of the 500 model neurons on a two-dimensional sheet, placed by a Halton quasi-random sequence; green, the 400 excitatory neurons; magenta, the 100 inhibitory neurons. **(b)** the membrane equation and the added ephaptic-field term, together with a diagram of the field (red) linking a neuron to its neighbors. The field on each neuron is the sum, scaled by the coupling strength, of its neighbors’ departures from rest. **(c)** normalized spatial coupling kernels versus inter-neuron distance: the ephaptic-field kernel (red) and the correlation of the input (grey). **(d)** the subthreshold voltage gain applied to each spatial mode of the population, in cycles per unit length: the closed-form transfer function H(k) (blue line), the field-coupled network (eLIF, red circles) and the field-free network (LIF, black squares). **(e)** Total spikes per trial for LIF (grey), eLIF (red) and RM (green). **(f)** pairwise subthreshold voltage correlation as a function of the distance between neuron pairs; the grey dotted line shows the correlation of the shared input. RM stands for rate-matched control. Error bars show the standard error of the mean (SEM); dots show individual network seeds. Asterisks show significance level: *p<0.05, **p<0.01, ***p<0.001.

To understand how this coupling would act on a whole population, we characterized the field first in space and then in spatial frequency. Neighboring neurons receive correlated inputs. Both the coupling and input correlation have Gaussian kernels falling off by distance, with the coupling kernel falling off over a shorter distance than the spatial correlation of the shared synaptic input (**Figure 1c**, **Figure S1b**). Thus, a neuron’s field is set mainly by its nearest neighbors. The kernel shape alone does not reveal what the field does to a spatially patterned population, because the field feeds back on the same activity that generates it. To make that effect explicit we computed the field’s transfer function, which gives the gain it applies to each spatial mode of the membrane potential and turns the local coupling rule into a prediction for the whole population. The field acts as a low-pass filter which amplifies the coarsest, near-uniform modes and leaves fine spatial structure untouched. A closed-form expression for the transfer function matched the simulated gain across modes almost exactly (**Figure 1d**; R-squared = .99), with the uniform mode amplified by 1.25 and the gain returning to unity by the third mode, and it held for a second kernel shape (**Figure S1c**). The field therefore reshapes the common, low-frequency component of population activity and leaves local structure alone. This common component is the shared drive that couples the trial-to-trial variability of nearby neurons, so it is the natural route by which the field could reshape population coding, and its consequences are what the rest of the paper tests.

We ran both the LIF and eLIF models across different seeds; the two are identical except for the ephaptic field term (see Methods). The ephaptic field elevated excitability in the network and raised the mean firing rate by 5.8% relative to LIF (**Figure 1e**; Wilcoxon signed-rank, p < .001, n = 12 seeds). This rise in rate is itself a consequence of the ephaptic model, but because a change in firing rate can by itself alter correlations and coding, and we wanted any effect to reflect the field rather than a change in excitability, we introduced a rate-matched control (labeled RM in the figures), the eLIF network with its baseline drive lowered until its firing rate returned to the LIF level (**Figure 1e**; matched in the conservative direction of firing slightly less than LIF, see Methods). Any residual difference between rate-matched control and LIF is then attributable to how the field reorganizes activity, not to a mere change in the network excitation. We first measured membrane-potential correlations to examine how coupling affects subthreshold covariability. Overall, the endogenous field reduced pairwise subthreshold voltage correlations at every distance (**Figure 1f**; from 0.20 to 0.05), and a smaller but reliable part of the reduction survived rate-matching (rate-matched control, from 0.20 to 0.17; Wilcoxon signed-rank, p < .001), which shows that the resting-state decorrelation is carried partly by the field and partly by the accompanying rise in firing rate. The reduction was evident and fairly uniform across distance, consistent with suppression of a broad shared mode rather than of local coupling, and it persisted across coupling strengths (**Figure S1d**) and across the grid of field and input length scales (**Figure S1e**).

### Endogenous fields decorrelate, imposed fields entrain

Ephaptic coupling and field effects on neurons have most often been studied and measured by imposing a field from outside the tissue, which paces and synchronizes firing (Jefferys, 1995; Frohlich and McCormick, 2010; Anastassiou et al., 2011). We therefore asked whether the way the field is delivered, that is either exogenously or endogenously, changes its consequences for coding. We swept the coupling strength from zero to near-critical and compared, under one identical field geometry, the endogenous eLIF field, its rate-matched control, and an exogenous field of the same spatial profile imposed open-loop from a fixed external source (**Figure 2a and 2b**; injected amplitude and distance attenuation in **Figure S2a and S2b**).

**Figure 2.**
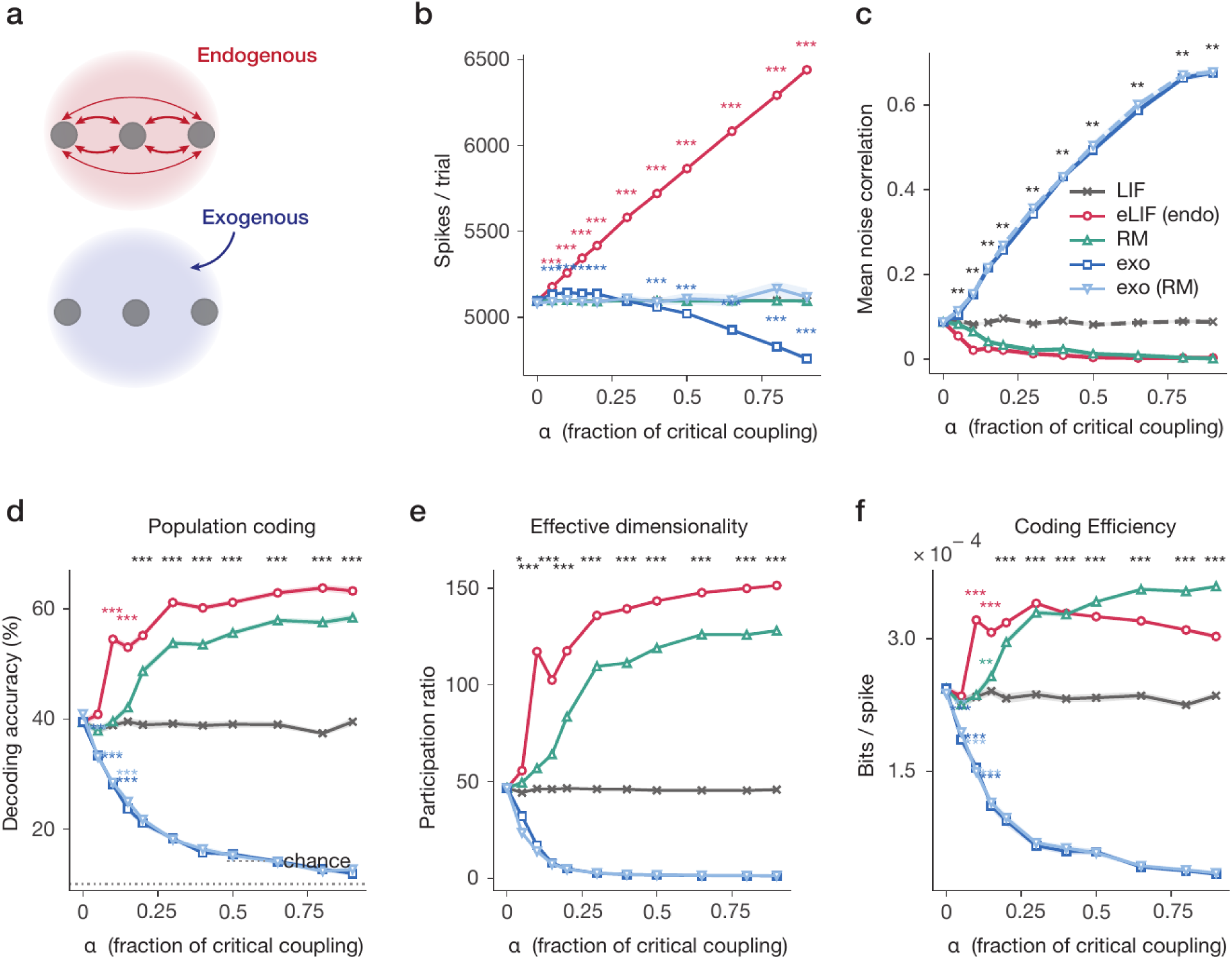
Comparison of endogenous and exogenous field effect on population coding. **(a)** Schematic of an endogenous field generated by the network activity (top) and an exogenous field imposed from an external source (bottom). **(b-f)** Network measures versus coupling strength α, expressed as a fraction of the critical value showing **(b)** Total spikes per trial, **(c)** Mean pairwise noise correlation, **(d)** Stimulus decoding accuracy (%), **(e)** effective dimensionality of the population response (the participation ratio), and **(f)** coding efficiency (bits per spike). RM stands for rate-matched control. Shaded error bars indicate the standard error of the mean (SEM); Asterisks show significance level: *p<0.05, **p<0.01, ***p<0.001.

These two field schemes moved population activity in opposite directions, with the endogenous field increasing and the exogenous field decreasing the firing rate (**Figure 2b**). The endogenous field reduced pairwise spike-count noise correlations (**Figure 2c**; from 0.09 to 0.004), whereas the imposed field elevated the noise correlation roughly sevenfold, to 0.70 (endogenous versus exogenous at the strongest coupling, Wilcoxon signed-rank, p < .001, n = 15 seeds); this opposite movement held in every network at both weak and strong coupling (**Figure S2c**).

To ask whether these opposite correlation structures mattered for coding, we characterized the population code in three complementary ways: the read-out accuracy for the stimulus (decoding), the effective dimensionality of the response (dimensionality), and the amount of information each spike carries (efficiency). We implemented a stimulus-coding task with ten broadly-tuned stimuli. All three measures separated the two schemes. Population decoding of stimulus identity rose from 39% to 63% under the endogenous field but fell to near chance under the imposed field (**Figure 2d**; 12%, chance = 10%; all p < .001; per-network in **Figure S2d**). The effective dimensionality, measured as the participation ratio of the response covariance spectrum (the number of dimensions the activity effectively occupies), expanded more than threefold for the endogenous field, from 47 to 152, but collapsed to about 1.2 under the imposed field (**Figure 2e**; per-network in **Figure S2e**; Wilcoxon signed-rank, p < .001, n = 15 seeds). Coding efficiency in bits per spike improved for the endogenous field and was cut to a fraction of baseline for the imposed one (**Figure 2f**; per-network in **Figure S2f**; Wilcoxon signed-rank, < .001, n = 15 seeds). Every gain survived rate-matching, for both the endogenous and the imposed field, showing that the effects were not driven solely by changes in population excitability and firing rate. In particular, the rate-matched endogenous model rose monotonically with coupling and tracked eLIF across all four measures (Spearman ρ = .97), and because it spends no extra spikes its per-spike efficiency gain exceeded that of every other model as coupling grew (**Figure 2f**). The benefit is therefore a result of the closed loop rather than of the field’s shape or of a change in rate as the same shape imposed open-loop is actively destructive to population coding and efficiency.

### Field coupling improves stimulus decoding

To connect these population-level changes to a defined computation, we trained the network on a ten-way discrimination of stimulus phase and read the stimulus identity from the population activity (**Figure 3a**; rate control and example rasters in **Figure S3a and S3b**). We fixed the coupling at one fifth of critical (alpha = 0.2), a value in the lower, biophysically conservative part of this range, so that the coding results do not depend on operating near the critical point. The field did not sharpen single-cell selectivity. Tuning curves were broad and essentially unchanged across models, so any improvement in population coding had to arise from the structure of trial-to-trial variability rather than from the cells’ tuning (**Figure 3b**; per-cell tuning correlation between LIF and eLIF, median r = .85, n = 400 cells, see the population tuning map, the example-cell tuning, and the coverage of stimulus space in **Figure S3c-e**).

**Figure 3.**
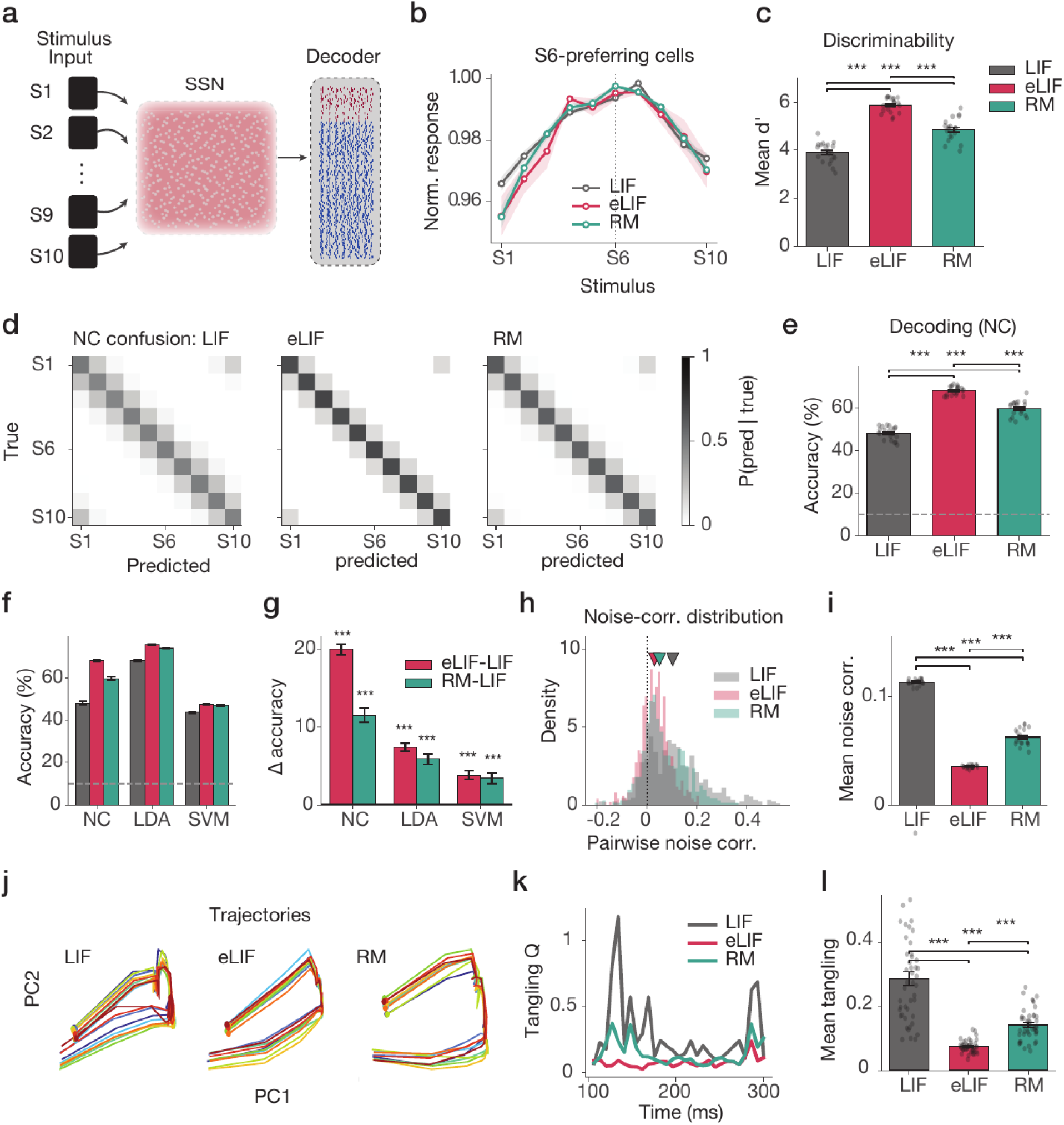
Stimulus decoding by the field-coupled network. **(a)** Task schematic: each trial one of ten stimuli (S1 to S10) activates the spiking network (SSN), and a decoder reads the stimulus response from the population spike counts. **(b)** Normalized tuning curves of the cells that prefer stimulus S6, versus stimulus identity **(c)** Mean neural discriminability (d′) between the stimulus responses, for the three networks. **(d)** Nearest-centroid (NC) confusion matrices for LIF, eLIF and RM. **(e)** Mean NC decoding accuracy (%) for the three networks; the dashed line is chance (10%). **(f)** Decoding accuracy (%) for three decoders (NC, LDA and SVM), each shown for the three networks; the dashed line is chance. **(g)** The change in decoding accuracy relative to LIF for eLIF (red) and RM (green), for each decoder. **(h)** The distribution of pairwise noise correlations across neuron pairs for for the three networks; triangles mark the medians. **(i)** Mean pairwise noise correlation for the three networks. **(j)** Population activity projected onto its first two principal components (PC1 and PC2) for LIF, eLIF and RM; each colored line is the trajectory for one stimulus. **(k)** Trajectory tangling (Q) over time for the three networks. **(l)** Mean trajectory tangling for the three networks. SSN stands for spiking neural network; RM, rate-matched control; d′, discriminability index; NC, nearest-centroid; LDA, linear discriminant analysis; SVM, support-vector machine; PC, principal component. Error bars show the standard error of the mean (SEM); dots show individual network seeds. Asterisks show significance level: *p<0.05, **p<0.01, ***p<0.001.

Field coupling improved neural discriminability, a measure of how far apart the stimulus responses sit relative to their trial-to-trial spread, from a d-prime of 3.9 to 5.9 for eLIF and 4.9 for the rate-matched control (**Figure 3c**). Consistent with this elevated discriminability, nearest-centroid decoding accuracy increased under the endogenous field, from 48% for LIF to 68% for eLIF and 60% for the rate-matched model (**Figure 3d and 3e**; Wilcoxon signed-rank, p < .001, n = 20 seeds). We examined other classifiers, including linear discriminant analysis (LDA) and support-vector machine (SVM) decoders, and the advantage held for these as well (**Figure 3f and 3g**; confusion matrices in **Figure S3f and S3g**). The gain was largest for the nearest-centroid read-out (20 points) and smaller for the more powerful decoders (7 points for LDA, 4 for SVM), yet it remained significant for every decoder. The extra structure the field adds is therefore genuinely decodable information rather than a crutch for a weak read-out.

To examine the population factors behind this read-out improvement, we asked whether the field reorganizes population variability. First, the field reduced pairwise noise correlations across the population (**Figure 3h and 3i**; mean from 0.11 to 0.035, a reduction of 69% for eLIF and 45% for rate-matched control; p < .001, n = 20 seeds), suggesting that it removes the shared, correlated component of trial-to-trial variability that would otherwise limit the population code. We also asked whether the field changes the geometry of the population dynamics in a way that benefits decoding. The low-dimensional population trajectories showed less crossing and overlap than under LIF (**Figure 3j**). Trajectory tangling, a measure of how often nearby population states move in different directions, the moments a downstream reader is most likely to confuse (Russo et al., 2018), was reduced for both eLIF and rate-matched control, indicating that the trajectories for different stimuli were less tangled (**Figure 3k and 3l**; mean from 0.29 to 0.075 for eLIF and 0.14 for rate-matched control; eLIF versus LIF Mann-Whitney U = 17, both p < .001). These effects persisted at matched rate, consistent with a coding benefit that reflects reduced shared variability (Averbeck et al., 2006; Cohen and Kohn, 2011) and that only partly depends on the field’s small increase in rate.

### Field coupling improves sparse temporal coding

To test whether the advantage regarding population coding generalized to other tasks, we examined a second task that involved temporal rather than spatial coding. The network had to classify four temporal activation patterns, an early decay, a late rise, a center peak, and a sustained drive, each delivered to a small, randomly chosen subset of neurons, while we varied both the sparsity of activation and the level of input noise (**Figure 4a**). Overall, accuracy increased with the number of active neurons and fell with noise, with eLIF above rate-matched control above LIF at essentially every point on the grid (**Figure 4b**; slices across sparsity and across noise in **Figure S4a**).

**Figure 4.**
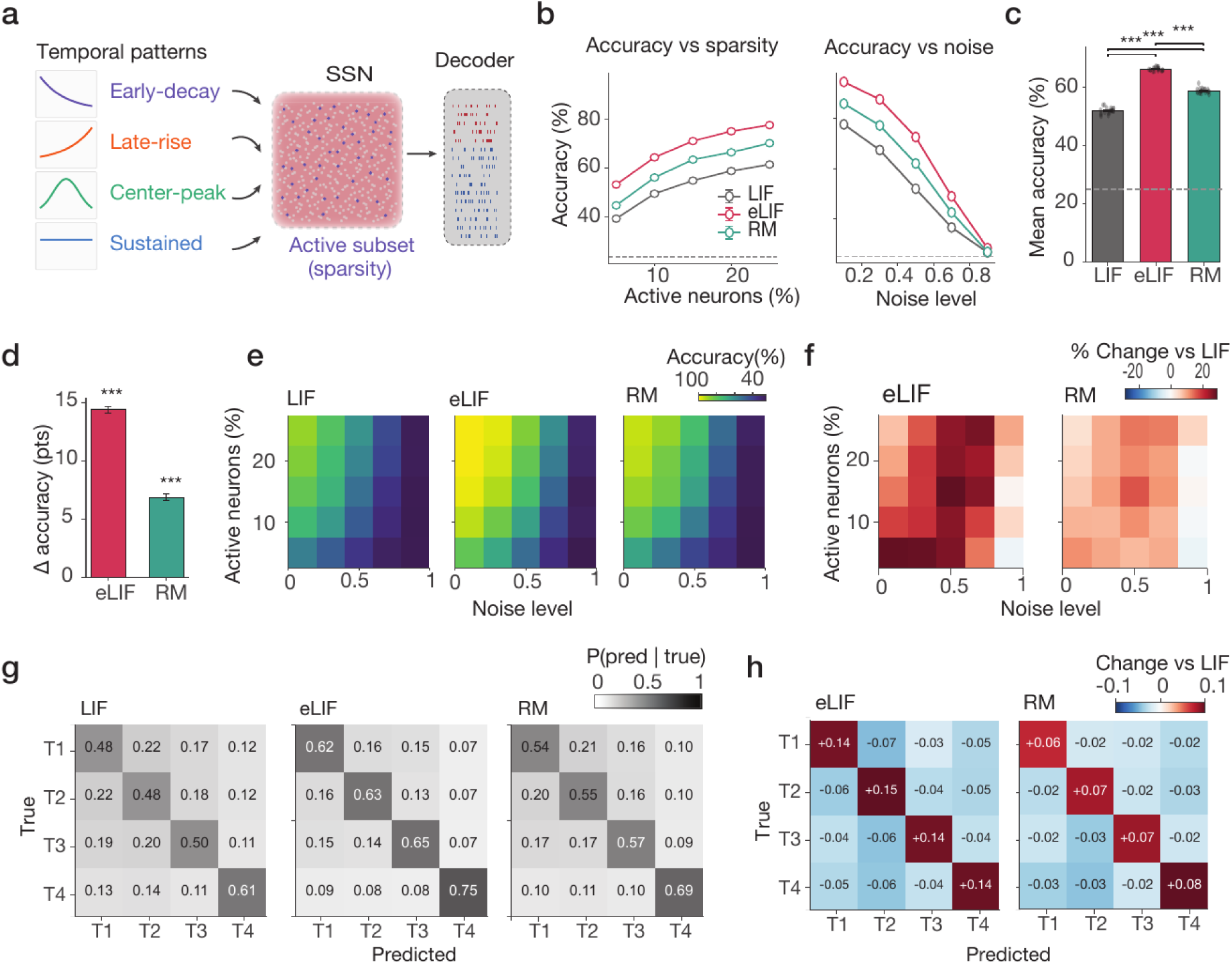
Sparse temporal-pattern classification. **(a)** Task schematic: one of four temporal activation patterns (early-decay, late-rise, center-peak or sustained) activates a subset of neurons (the sparsity) in the spiking network under variable noise conditions, and a decoder classifies the pattern. **(b)** Classification accuracy as a function fraction of active neurons (left) and the input noise level (right), for the three networks. **(c)** Mean accuracy averaged across all the sparsity and noise levels for the three networks; the dashed line is chance (25%). **(d)** The change in accuracy relative to LIF, in percentage points, for eLIF (red) and RM (green). **(e)** Heatmap accuracy across the grid of the sparsity versus noise level. **(f)** The percentage change in accuracy relative to LIF across the same grid, for eLIF and RM. **(g)** confusion matrices for the four patterns (T1 to T4) in the three networks. **(h)** The difference in confusion entries relative to LIF, for eLIF and RM. T1 to T4 denotes the four temporal patterns; LIF, eLIF and RM as defined in Figure 1. RM stands for rate-matched control. Error bars show the standard error of the mean (SEM); dots show individual network seeds. Asterisks show significance level: *p<0.05, **p<0.01, ***p<0.001.

Averaged over the sparsity-by-noise grid, nearest-centroid accuracy was 52% for LIF, 66% for eLIF, and 59% for rate-matched control, all well above the 25% chance level (**Figure 4c**; per-seed spread in **Figure S4b**). The field improved accuracy by 14.4 points, of which 6.9 points, about half, survived rate-matching (**Figure 4d**; Wilcoxon signed-rank, p < .001, n = 20 seeds). The gain was present throughout the grid except at the highest noise level, where it fell to 1.7 points (**Figure 4e and 4f**). The confusion matrices show that the improvement was a uniform strengthening of the correct-class rate across all four patterns rather than a repair of a single confusable pair, with every diagonal entry increasing and every off-diagonal entry decreasing (**Figure 4g and 4h**; a linear decoder gives the same ordering, **Figure S4c**). The rate-matched benefit also generalizes across sparsity and across low to moderate noise. Together, these results show that field coupling improves the population code both for stimulus identity and for the temporal structure of the input.

### A field signature in cortical tissue

If cortical neurons are ephaptically coupled, one cell’s spike should leave a small, time-locked trace in the subthreshold voltage of a nearby cell, and that trace should weaken with distance. We probed this signature in the Allen Institute synaptic physiology dataset, in which up to eight neurons were patched simultaneously (**Figure 5a and 5b**; Campagnola et al., 2022). For every ordered pair of unconnected neurons, we averaged one cell’s subthreshold voltage on the spikes of another, using only isolated spikes with no other recorded cell firing within 15 ms and excluding synaptically connected pairs (69 experiments, 928 unconnected pairs).

**Figure 5.**
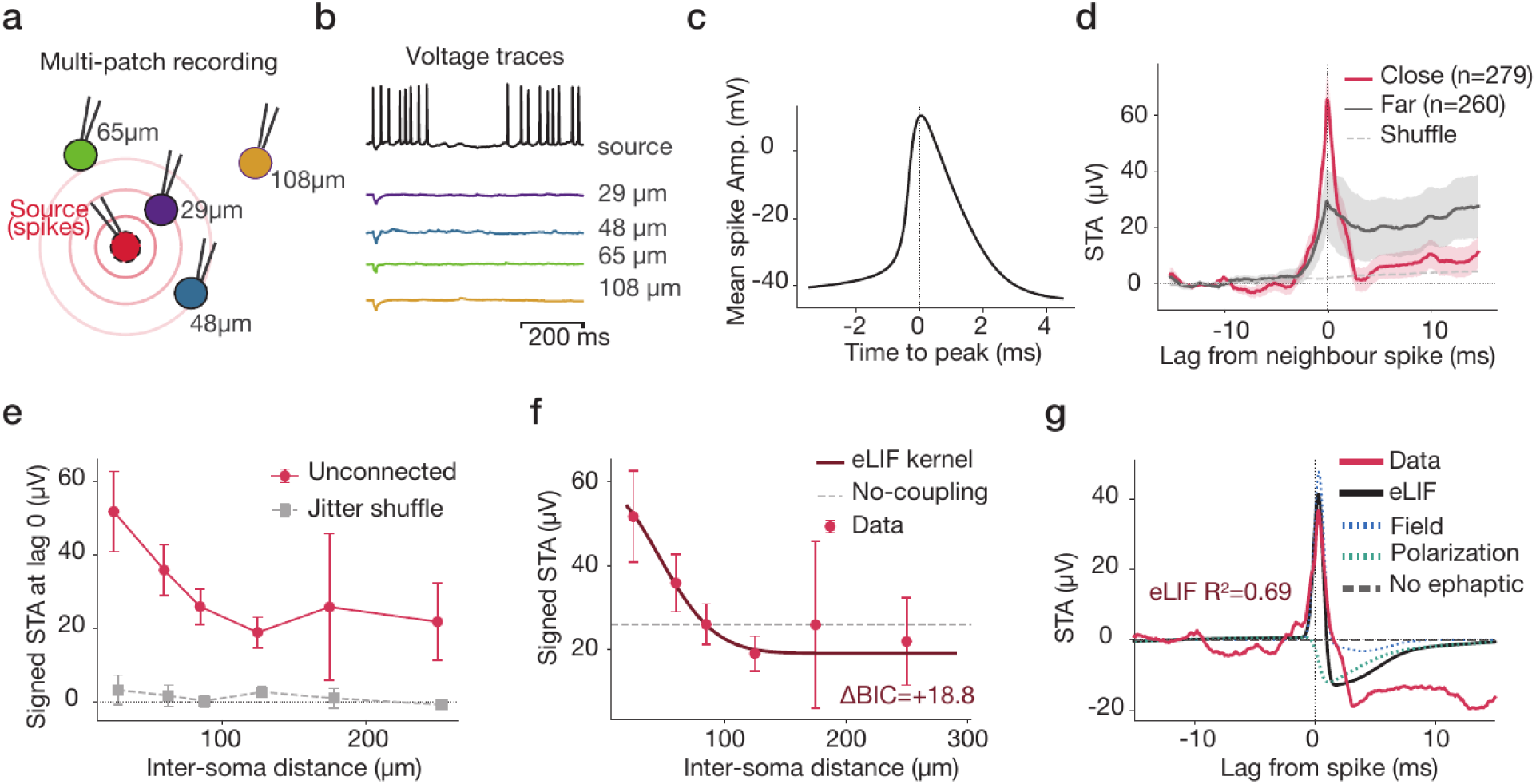
A spike-triggered subthreshold signature in cortical multi-patch recordings. **(a)** Schematic of a multi-patch experiment: spikes from a source neuron are related to the subthreshold voltage of simultaneously recorded neurons at different inter-soma distances. **(b)** Example simultaneous voltage traces from the source neuron and four neighbors. **(c)** The mean action-potential waveform of an example source neuron. **(d)** The spike-triggered average (STA) of a neighbor’s voltage as a function of lag from the source spike, for close pairs (less than 70 µm), far pairs (more than 150 µm) and a jitter-shuffled control. **(e)** The signed STA amplitude at zero lag as a function of inter-soma distance, for unconnected pairs (red) and the jitter-shuffle control (grey). **(f)** The signed STA amplitude with the eLIF field-kernel fit (black) and the no-coupling model (dashed); compared by the change in Bayesian information criterion (ΔBIC). **(g)** The full time course of the distance-dependent component (the excess STA of close minus far pairs), with the eLIF model fit and its two components (the instantaneous field and the membrane polarization) and the non-ephaptic prediction. Error bars indicate the standard error of the mean (SEM).

The triggering event was a stereotyped action potential (**Figure 5c**), and the spike-triggered average of each neighbor’s membrane potential was computed for every pair. Spikes were followed by a small deflection in the neighbor’s membrane potential, time-locked to the spike and rising above a jitter-shuffled null (**Figure 5d**). As a positive control, connected pairs showed the expected larger and slower postsynaptic waveform, distinct from the small deflection seen in unconnected pairs (**Figure S5a**). The deflection was larger for nearby pairs. The signed amplitude was significantly larger for close pairs (< 70 µm) than for far pairs (> 150 µm) (**Figure 5d**; Mann-Whitney, p < .001, n = 279 close and 260 far pairs). This close-minus-far gap was present in most individual experiments and across distance bins (**Figure S5b and S5c**), and the deflection declined with inter-soma distance across the population (**Figure 5e**; Spearman ρ = −.21, p < .001, n = 928; per-pair values over the full sampled distance range in **Figure S5d and S5e**). The distance profile was fit by the eLIF field kernel, which a model comparison favored over a no-coupling model (**Figure 5f**; delta BIC = +18.8). The full time course of the distance-dependent component, a fast positive transient near the spike followed by a slow polarization lobe, was reproduced by the eLIF model that combines the instantaneous field with the neighbor’s membrane filtering (**Figure 5g**; R^2^ = .69; full lag-by-distance average in **Figure S5f**), whereas the non-coupled model predicts no deflection.

Together, these recordings reveal a small but spike-locked and distance-dependent subthreshold deflection between unconnected cortical neurons, matched in amplitude and time course by the same field model that produced the coding effects in simulation, as expected if cortical tissue is weakly ephaptically coupled.

## Discussion

We asked whether a self-generated electric field shapes the neural population code, and whether a closed-loop field differs from one imposed from outside. We found that, treated as a self-consistent feedback term, the field behaves as a low-pass filter that acts on the coarsest, shared mode of the population and leaves fine structure untouched (**Figure 1**). Importantly, the way the field is integrated into the network can reverse its effect. An endogenous field reduces noise correlations, expands the dimensionality of the population response, and raises decoding accuracy and coding efficiency, whereas the identical field imposed open-loop moves every measure the other way and reduces coding accuracy (**Figure 2**). The endogenous benefit is a reorganization of population variability rather than of rate. It survives matching the firing rate to the field-free network, raising discriminability and decoding while lowering noise correlations and trajectory tangling, and it holds both in a stimulus-identity decoding task and in a sparse temporal-classification task (**Figure 3 and Figure 4**). The field model that produces these effects predicts a small, spike-locked, distance-dependent subthreshold deflection between unconnected neurons, which we detect in cortical multi-patch recordings (**Figure 5**).

The opposite effects of endogenous and exogenous field integration (**Figure 2**) help address an existing gap in the literature. Fields imposed from outside typically pace and synchronize neurons, from single cells to networks (Deans et al., 2007; Ozen et al., 2010; Krause et al., 2019). Even the studies framed around the endogenous field test it by applying a field from an electrode, matched to or modulated by the ongoing activity, rather than isolating the intrinsic one, which cannot be manipulated without changing the activity that produces it (Frohlich and McCormick, 2010; Reato et al., 2010; Anastassiou and Koch, 2015). A genuinely self-generated field has been shown to act on its own mainly in the non-synaptic propagation of slow or epileptiform waves, where activity recruits its neighbors through the field it makes (Zhang et al., 2014; Qiu et al., 2015; Chiang et al., 2019), but that is a spreading of synchrony rather than a test of a population code or of how activity organizes its geometry moment to moment. Our results place the difference in the loop rather than in the field. An externally imposed field is a low-dimensional drive common to all neurons and applied from outside, and in our network it collapses the response onto a low-dimensional synchronized subspace, as the entrainment literature predicts (Deans et al., 2007; Reato et al., 2010). The same field, generated by the activity it acts on and returned to it, instead reduces the shared variability and expands the dimensionality of the code. The way the field is integrated into the network, and not its spatial shape, therefore sets the sign of the effect, which cautions against reading the function of the endogenous field directly from open-loop stimulation.

Another finding is that ephaptic coupling can also be seen as an operation on the population code, not on rate or synchrony alone. Most functional studies of fields have asked whether they shift spike timing or entrain rhythms (Radman et al., 2007; Reato et al., 2013); we ask instead what they do to the shared variability that can limit how much a population encodes (Averbeck et al., 2006; Cohen and Kohn, 2011). In the model the endogenous field removes much of this shared component, lowering pairwise noise correlations, expanding the number of dimensions the response occupies, and untangling the trajectories a downstream reader has to separate (**Figure 2 and Figure 3**; Russo et al., 2018). We do not claim that the field adds information along the stimulus-limiting dimension in the strict sense of Moreno-Bote et al. (2014); rather, it reshapes the correlation structure into a form that a fixed decoder can exploit (**Figure 3**). This suggests that the self-generated field acts as an active organizer of population geometry, a complement to the recent proposal that the cortical field also carries information of its own (Pinotsis and Miller, 2022, 2023).

A main difficulty in this area is that any consequence of the endogenous field co-varies with the activity that produces it (Anastassiou and Koch, 2015). Our rate-matched control addresses this directly, by lowering the baseline input until the field-coupled network fires at the field-free rate, which separates the field’s reorganization of activity from the change in excitability it brings. We showed that about half of each coding effect remains at matched rate (**Figure 3 and Figure 4**), which indicates that a substantial part of the benefit is carried by the structure the field imposes on variability rather than by the extra spikes it elicits. Keeping the field fixed while changing the rate has no direct experimental counterpart, but it lets the model separate the structural effect from the change in excitability.

While the recordings complement our computational findings, they are meant as support rather than proof. We showed, in the Allen Institute multi-patch dataset (Campagnola et al., 2022), that one neuron’s subthreshold voltage exhibits a small deflection time-locked to the spikes of an unconnected neighbor, larger for nearby pairs and declining with distance, and reproduced in both amplitude and time course by the same field kernel used in the model (**Figure 5**). The effect is on the order of tens of microvolts and its distance dependence is weak, so we read it as consistent with, not proof of, ephaptic coupling. Residual electrode cross-talk cannot be fully excluded, but its lag structure and distance dependence argue against it (**Figure S5**). Two features nonetheless make it useful. First, its amplitude places cortical tissue at the low-coupling end of the range we simulated, below the point at which the modeled effects are largest, so the coding benefits we report do not require the network to sit near criticality (**Figure 2**). Second, its distance profile is better explained by the same field-kernel form used in the model than by a no-coupling account, which ties the simulation and the recording together. A deflection of tens of microvolts from a single neighbor’s spike is consistent with the sub-millivolt somatic polarizations that whole physiological fields, summing many active neurons, produce in cortex (Anastassiou et al., 2011; Anastassiou and Koch, 2015).

In summary, a self-generated field acts as a spatial filter that reorganizes the shared, low-frequency component of population activity and, in doing so, improves the code the population carries. It explains two observations that have been hard to unify; externally imposed fields synchronize while a self-generated field does not, and a coupling too weak to change firing much can still change what the activity represents. It also makes testable predictions, though they turn on the field being internally generated rather than merely present. Because an imposed field of the correct spatial profile still synchronizes in our model, an externally applied field, including closed-loop stimulation controlled by the recorded activity, should entrain rather than decorrelate; the decorrelating effect should instead appear when the intrinsic coupling is strengthened, for example by raising the extracellular resistivity, which scales the field a given current produces (Buzsaki et al., 2012). Such a change should lower noise correlations and expand dimensionality, with the benefit tracking the shared rather than the private component of variability. Whether cortex uses this channel is therefore still open, but our results indicate that if neurons are coupled by their shared field at all, even weakly, that field is positioned to help rather than harm the population code.

## Methods

We used spiking neural network (SNN) modeling as the main approach to simulate how ephaptic (electric-field) coupling might contribute to population coding and its functional roles in cortical neural networks. We first examined the effect of endogenous versus exogenous field on population dynamics, then continued with the endogenous field for the rest of the paper. Finally, we show that such modeling of field effect endogenously is biologically realistic.

### Network model

#### Single-neuron dynamics

Each neuron was modeled by a leaky integrate-and-fire (LIF) spiking neuron model, which is a standard reduction of a neuron to a single membrane capacitor that leaks charge and generates a spike when its voltage crosses a threshold (Lapicque, 1907; Tuckwell, 1988; Burkitt, 2006; Gerstner et al., 2014). For each neuron the membrane potential *V_i_* then obeys

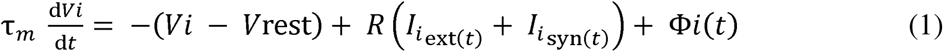

where τ_m_ is the membrane time constant, rest is the resting potential, and R= 100 MΩ is the membrane (input) resistance. is the external current driving neuron i (baseline, noise and stimulus) and is the recurrent synaptic current. The term Φ_i_(t) is the ephaptic (electric field) contribution (explained in Ephaptic field coupling section), reflecting endogenous field from the activity of the nearby neurons, which is present for modeling the ephaptic coupling and set to zero for the standard model in the absence of any ephaptic coupling effects. When *V_i_* reaches the threshold *V_th_*, the neuron generates a spike, and the membrane voltage is reset to *V*_reset_ and held there for an absolute refractory period τ_ref_:

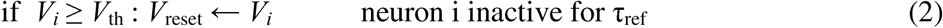

All parameter values were chosen within the ranges measured for neocortical pyramidal neurons and literature: τ_m_ = 10 ms, *V*_rest_ = −60 mV, *V_th_* = −50 mV, *V*_reset_ = −60 mV (reset to rest) and τ_ref_ = 2 ms (McCormick et al., 1985; Markram et al., 2015). We implemented these differential equations using the forward Euler method with a fixed time step Δt = 0.1 ms.

### Synaptic coupling

Synaptic currents in eq. 1 were modeled with single-exponential functions, a computationally efficient description of synaptic transmission (Destexhe et al., 1998; Gerstner et al., 2014). Synaptic current to each neuron is computed by the summation of total excitatory and inhibitory currents. The recurrent synaptic current of neuron i is then modeled as:

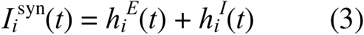

where each current was elicited and decayed by firing of presynaptic neurons by:

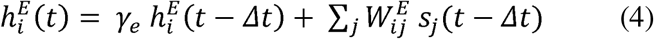

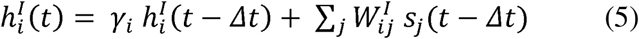

Here, 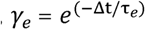, and 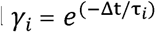 are decay factors with τ_e_ = 3 ms and τ_i_ = 8 ms excitatory and inhibitory synaptic time constants, respectively (Salin and Prince, 1996; Destexhe et al., 1998). s_j_(t) is a binary variable for spike train of neuron j, and are excitatory and inhibitory synaptic weights from neuron j onto neuron i, and synaptic weights are drawn from a Gaussian distribution with a mean effective voltage of 0.4 mV and standard deviation 0.1 mV for excitatory, and a mean −1.5 mV and standard deviation of 0.3 mV for inhibitory synapses (Markram et al., 2004).

### Network architecture and geometry

The network contained N = 500 neurons, with N_E_ = 400 (80%) excitatory and N_I_ = 100 (20%) inhibitory neurons, consistent with the approximate ratio observed and modeled in the cortex (Markram et al., 2004; Banaie Boroujeni et al., 2023, 2025). Each neuron was assigned a fixed position on the unit square by a Halton quasi-random sequence with bases 2 and 3 (Halton, 1964) with spatial coordinates in normalized units of the square side. Excitatory and inhibitory cells were mixed and their connectivity was random (Erdos-Renyi) with a connection probability of p = 0.15 (Holmgren et al., 2003; Perin et al., 2011). The connectivity graph was fixed by a separate random seed so that simulations on all neuron models within a run shared identical wiring.

### External drive and noise

Each neuron was fed with an external input which was the sum of a constant baseline drive, a private noise, a spatially structured shared-noise, and a stimulus-driven current input implementing task specific activation of each neuron. Private noise reflects cell-specific input fluctuations and is modeled by an independent Gaussian white noise added to each neuron. Shared noise was generated by K = 50 noise sources randomly placed in the network (K is picked to be large enough to generate a smooth spatial structure yet far smaller than N). Each source is a smoothened independent white-noise signal distributed to neurons through a distance-weighted, row-normalized mixing matrix G:

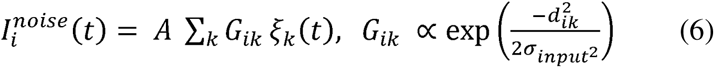

Here G_ik_ is the weight of noise source k on neuron i, A is the shared-noise amplitude, and σ_input_ = 0.15 is a spatial scale for shared noise correlation. This generates a correlated noise between nearby neurons with no task information, which is a simple model of shared input fluctuations within spiking networks (Doiron et al., 2016).

### Ephaptic field coupling

We developed an ephaptic LIF (eLIF) neuron model using a term reflecting the local extracellular field generated by the weighted average of transmembrane potential of nearby neurons (Jefferys, 1995; Anastassiou and Koch, 2015; Goldwyn and Rinzel, 2016). This ephaptic term reflects an effective depolarization, proportional to a spatially weighted sum of neighboring membrane deflections from rest and added to the neuron model by Φ*_i_*:

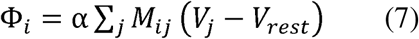

where V_j_ - V_rest_ is the membrane potential deflection of neuron j from rest, M_ij_ is the spatial coupling weight from neuron j onto neuron i which constitutes a row-normalized Gaussian kernel of neighboring activity seen at each neuron, and α is a scalar indicating coupling strength defined below.

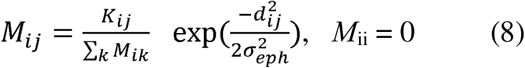

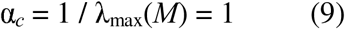

where K_ij_ is the unnormalized Gaussian weight between neurons i and j, d_ij_ denotes the Euclidean distance between neurons i and j, the field length scale σ_eph_= 0.15 sets how far the field reaches. Coulomb-type (softened point-source) and global mean-field kernels were also implemented. Every kernel was scaled so that its largest eigenvalue λ_max_(*M*) equals one, so α is the fraction of the critical coupling at which the linear response of the network diverges (default α = 0.2, that is 20% of critical).

### Neuron models: standard LIF, eLIF, and rate-matched

Three models were compared throughout. The standard LIF model, hereafter LIF, is the network with the field term removed (Φ_i_ = 0). The eLIF model adds the field term with all other parameters and all random inputs held identical to LIF, so that any difference between LIF and eLIF is attributable to the field alone. Because the field can change the excitation level in the network and firing rate, we added a rate-matched control (labeled RM in the figures). Rate-matched control is the eLIF model with a uniform constant offset subtracted from the baseline drive so that its spike count in the stimulus window equals that of the LIF model. The offset was found by bisection through an iterative process on a short probe of the network and applied only when eLIF fired more than 1% above the LIF rate. The rate-matched control aims to separate the effect of the field on the population code from any change in firing rate that could be attributed to the excitability. For general comparisons between the networks, voltage correlations were computed from downsampled membrane traces in the stimulus window and binned by pairwise distance; the LIF and eLIF models were compared per distance bin by a two-sided Mann-Whitney U test. We also measured local field potential and its power spectrum (Welch periodogram, Hamming window, 0.1 to 100 Hz; Welch, 1967), the dependence of voltage correlation on the field width σ_eph_, the kernel structures, and the sensitivity of decoding and correlation to α (**Figure S1**).

### Linear response and spatial transfer function

To characterize the spatial action of the field and to obtain its critical coupling analytically, we computed the linear (subthreshold) response of the network, which in the spatial Fourier domain reduces to a single gain H(k) at each wavenumber k. The field couples selectively to coarse, spatially coherent modes (low k) and is nearly transparent to fine, private ones (high k); the low-mode gain 1/(1 - α) diverges as α approaches 1, which sets the critical coupling and is why α is reported as a fraction of critical.

In the subthreshold regime, the network is linear and its steady-state response to a constant input pattern is

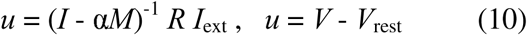

where u is the vector of steady-state membrane deflections from rest, I is the identity matrix, and I_ext_ is the external input pattern. Each spatial eigenmode m of the kernel, with eigenvalue λ_m_, is amplified by the gain 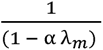. For a translation-invariant Gaussian kernel M is diagonal in the spatial Fourier basis, with eigenvalue 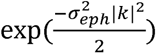 at spatial wavenumber k, so the spatial transfer function, the gain as a function of wavenumber, is

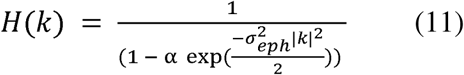

We tested this prediction by driving the network with spatial sinusoids (wavenumbers m = 0 to 12) with spiking disabled and comparing the measured subthreshold gain to the analytic H(k) (**Figure 1f**).

### Endogenous versus exogenous field coupling

To ask whether the field’s effect depends on how it is generated and not only on the operator, we examined coupling as endogenously and exogenously generated (**Figure 2**). In endogenous (closed-loop) coupling the field reads the real-time membrane voltages of the network itself, as in eq. 7; meaning that the field is a function of the network state and feeds back on it. In exogenous (open-loop) coupling an external field with the same spatial operator is imposed on the network but does not depend on the network state. We modeled the exogenous field as a single source at the center of the square whose influence decayed with distance and was modulated by a shared temporal envelope:

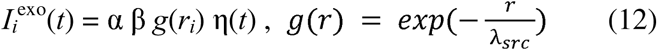

where r_i_ is the distance of neuron i from the source, g(r) is the spatial attenuation profile with decay length λ_src_ = 0.5, β = 8 mV per unit α is the amplitude scale, and η(t) is a unit-variance shared temporal envelope (white noise smoothed by the same 50 ms boxcar used for the network noise). The exogenous field is therefore a state-independent standing pattern shared across the population, the regime expected for an externally applied or strongly volume-conducted field.

Five conditions were compared as a function of α: standard (no field), endogenous, rate-matched endogenous, exogenous, and rate-matched exogenous. The sweep used α in {0, 0.1, 0.2, 0.3, 0.4, 0.5, 0.65} and twelve independent networks, with a fresh network and fresh random initial voltages drawn uniformly between reset and threshold for each seed; the same initial conditions were used across the five conditions within a seed. To characterize the population, we then compare mean pairwise noise correlation, population synchrony (defined below), and participation-ratio dimensionality. Endogenous and exogenous conditions were compared at each α by a paired Wilcoxon signed-rank test with Holm correction across the α values.

### Coupling strength as a control parameter

To characterize how the strength of the coupling affects the population code, we swept α as a control parameter from zero toward criticality, α in {0, 0.05, 0.1, 0.15, 0.2, 0.3, 0.4, 0.5, 0.65, 0.8, 0.9}, with fifteen independent networks per point (**Figure 2**). At each value of α the standard, eLIF and rate-matched models were run on the same freshly generated network, so the three models are paired. The standard model was recomputed at every α to account for seed-to-seed fluctuations. The task used ten broadly tuned stimuli (the decoding task, below) with fifty trials each. As a function of α we measured nearest-centroid and linear-discriminant decoding accuracy, mean noise correlation, population synchrony, participation-ratio dimensionality, coding efficiency in bits per spike, and spikes per trial. eLIF was compared to the standard model at each α by a paired Wilcoxon test with Holm correction across α (Figures 2,S3).

### Coding tasks, decoders, and population metrics

#### Stimulus decoding task

The decoding task asked how well a downstream reader could recover stimulus identity from a single trial of population activity (**Figure 3**). There were ten stimuli. Each excitatory neuron was assigned a random preferred stimulus with a circular Gaussian tuning curve over stimulus identity as:

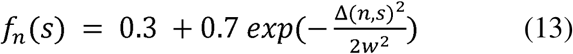

where f_n_(s) is the relative drive to neuron n for stimulus s, Δ(n, s) is the circular distance between the preferred stimulus of neuron n and the presented stimulus s, and w is the tuning width, set to w = 8. Each trial duration was 400 ms; with the stimulus on from 100 to 300 ms. The per-neuron input during the stimulus was a baseline of 10 mV, a shared broad drive of 2 mV to all excitatory cells, and a stimulus-specific drive of 2 mV scaled by the tuning curve, with shared and private noise amplitudes of 4 mV each (effective voltages R times I, as defined above). The signal was therefore a small modulation on a large noisy drive. Twenty independent networks were run with 150 trials per stimulus. The feature vector for each trial was the 500-dimensional vector of per-neuron spike counts in the count window.

### Sparse spatiotemporal coding task

The sparse task tested whether the field benefit generalizes to spatially sparse, temporally structured patterns (**Figure 4**). There were four patterns, each defined by a fixed random subset of excitatory cells (the spatial code) driven by one of four temporal profiles over an 80 ms window: an early-decay exponential, a late-rise exponential, a center-peaked Gaussian, and a sustained profile. Sparsity was defined by fraction of excitatory cells with pattern-specific external drive and swept over {0.05, 0.10, 0.15, 0.20, 0.25}. The noise level was swept over {0.1, 0.3, 0.5, 0.7, 0.9}. The pattern-specific drive scaled down with noise as drive proportional to (1 minus noise level), on top of a 10 mV baseline and a 2 mV shared drive to all excitatory cells, with shared and private noise of 4 mV. Each trial duration was 150 ms with the injection window from 35 to 115 ms. Twenty independent networks were run with 40 trials per pattern.

### Decoders

Decoders were trained and tested on the per-trial population spike-count vectors. Chance accuracy is one over the number of stimuli/patterns, that is 10% for the ten-stimulus decoding task and 25% for the four-pattern sparse task. We used three different scores for the decoder task to show the effect is independent of the classifier type, but largely used Nearest centroid (NC) for all other tasks and plots.

NC classifier used leave-one-out cross-validation where for each held-out trial the class template was computed from the mean vector of remaining trials, and the class assigned based on the nearest template in squared Euclidean distance:

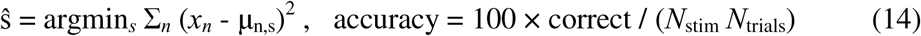

where x_n_ is the spike count of neuron n on the test trial, μ_n,s_ the mean template for neuron n and stimulus s computed over the training trials, ŝ is the predicted stimulus, and N_stim_ and N_trials_ are the numbers of stimuli and trials.

In addition to NC, we used linear discriminant analysis (LDA; Fisher, 1936) and a support vector machine (SVM; Cortes and Vapnik, 1995) with stratified five-fold cross-validation on z-scored features using scikit-learn (Pedregosa et al., 2011); and computed mean accuracy across folds. In stimulus-decoding task, we also used pairwise discriminability derived by the sensitivity index d-prime (Green and Swets, 1966). For two stimuli a and b,

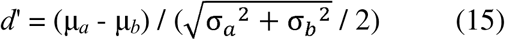

where μ_a_and μ_b_ and σ ^2^ and σ ^2^ are the mean and variances of the two stimuli responses, computed on the projection of the population counts onto the unit vector. The reported value was the mean over all distinct stimulus pairs. Coding efficiency was decoding accuracy divided by spikes per trial.

### Population and information metrics

To characterize population coding dynamics we used several metrics, including noise correlations, effective dimensionality, coding efficiency, population synchrony, and trajectory tangling. Noise correlations were the average across stimuli of the Pearson correlation, taken across trials, of the trial-to-trial residual spike counts of neuron pairs (a fixed random set of pairs). Effective dimensionality was the participation ratio of the neuron-by-neuron count covariance (Gao and Ganguli, 2015; Litwin-Kumar et al., 2017), defined by:

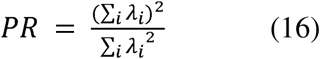

where λ_i_ are the eigenvalues of the covariance matrix; PR is small when activity is confined to a few dimensions and large when it spreads over many.

Stimulus information was defined by the Shannon mutual information between stimulus and decoded response, computed from the confusion matrix, in bits (Shannon, 1948; Quian Quiroga and Panzeri, 2009),

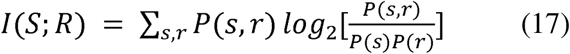

where S is the presented stimulus, R the decoded response, P(s, r) their joint probability, and P(s) and P(r) the marginals; coding efficiency in the bifurcation analysis was reported as bits per spike.

Population synchrony was measured on spike trains binned at 5 ms (Golomb and Rinzel, 1994),

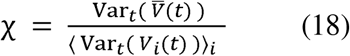

which is the ratio of the temporal variance of the population-averaged signal V (t) to the average over neurons of the single-cell temporal variances; χ approaches one for full synchrony and zero for asynchronous activity.

Population geometry was characterized by trajectory tangling on the first three principal components (PCs) (Russo et al., 2018),

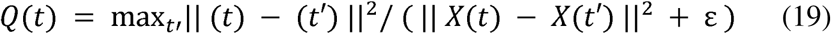

where X(t) is the population state at time t in PC space, is its time derivative, the maximum runs over all other times A, and ε = 0.1 is a small constant that prevents division by near-zero distances; high *Q* marks states where similar population activity moves in different directions. Low-dimensional trajectories were obtained by joint PC analysis of the trial-averaged, temporally smoothed population vectors of the standard and eLIF models.

### Noise structure and spatial frequency

To test when the field decorrelation actually helps coding (**Figure S3**), we decoded a weak coherent spatial cosine (eight phase classes) while varying the spatial correlation length σ of the noise, smoothing independent noise with a Gaussian of width σ over {0, 0.1, 0.2, 0.4, 0.6, 0.8, 1.0} at fixed total power, with one hundred independent networks per level at α = 0.2. We decoded stimulus phase for the standard, eLIF and rate-matched models and recorded the standard-model noise correlation. eLIF and rate-matched eLIF were each compared to the standard model by paired Wilcoxon tests with Holm correction across σ. A spatial-frequency control placed signal and noise at the same wavenumber. Wavenumbers m in {1, 2, 3, 4, 6, 8, 10, 12} were tested over five networks.

### Statistical analysis

Every figure point is a mean over independent network seeds, and error bars and bands are the standard error of the mean over those seeds (the standard deviation divided by the square root of the number of seeds). Each seed is a fully independent network with its own connectivity, positions and noise, so the seed is the unit of replication. Per-seed values were stored so that comparisons between models on the same seeds are paired. The numbers of seeds were set to twelve for the endogenous versus exogenous analysis (**Figure 2**), fifteen for the bifurcation sweep (**Figure 2**), one hundred for the noise-structure analysis (**Figure S3**), and twenty each for the decoding and sparse coding tasks (**Figures 3 and 4**).

When two models were compared on the same seeds we used the Wilcoxon signed-rank test (paired; Wilcoxon, 1945); when samples were unpaired or of unequal length we used the two-sided Mann-Whitney U test (Mann and Whitney, 1947). Per-seed differences compared with the standard model were tested against zero with the one-sample Wilcoxon test. Unless otherwise stated, tests were two-sided. Simulations and analyses were implemented in Python with NumPy (Harris et al., 2020) and SciPy (Virtanen et al., 2020).

Multiple comparisons were controlled by the Holm-Bonferroni step-down procedure (Holm, 1979), applied within each family of comparisons rather than globally. For the three-model figures the family was the three pairwise comparisons among standard, eLIF and rate-matched within each metric. For the sweep figures the family was the set of sweep points (across α for **Figure 2**, across σ for **Figure S3**).

### Real-data test of the ephaptic prediction

#### Dataset

We tested the central prediction of the model, a short-range post-spike voltage deflection in unconnected neighbors, on the Allen Institute for Brain Science Synaptic Physiology dataset, release 2.1 (Seeman et al., 2018; Campagnola et al., 2022). These are multi-patch whole-cell current-clamp recordings from acute cortical slices, with up to eight neurons recorded simultaneously. We used cell, pair and connectivity information, and the per-experiment Neurodata Without Borders (NWB) files for the raw membrane traces (Rubel et al., 2022). We used 71 experiments. Only recordings in current-clamp mode were analyzed, and membrane traces were handled in volts.

### Cells, pairs, and connectivity

Cells were mapped to recording electrodes through the cell and electrode tables of the database, and soma positions were read from the cell records. The inter-soma distance for each pair was converted to micrometers from the table. We labeled each pair as “connected” if it had either a chemical synapse or an electrical (gap-junction) connection, as recorded in the database; or “unconnected” otherwise. The minimum number of spikes for spike-triggered averages between pairs was 50 which yielded 999 ordered pairs (a pair contributes two ordered measurements, one in each direction), of which 928 were unconnected and 71 connected. Inter-soma distances for unconnected pairs ranged from 11 to 1430 micrometers, with a median of 101 micrometers.

### Spike detection and isolation

Spikes were detected as upward crossings of a −20 mV threshold, with crossings within a 2 ms refractory window merged. To remove contamination by coincident firing, a source spike was used only if it was isolated, meaning that no other simultaneously recorded cell spiked within 15 ms of it. Isolation was tested against the merged spike train of all other recorded cells in the same sweep.

### Spike-triggered average and signed deflection

For each ordered pair (source A, neighbor B) we computed the spike-triggered average of the neighbor membrane potential (±15 ms), with the mean of the neighbor trace in a pre-spike window from −15 to −8 ms subtracted from each segment. The reported signed deflection was the mean of the spike-triggered average over the interval within ±1 ms of the source spike. Pairs were grouped by distance into close (below 70 micrometers) and far (above 150 micrometers). Several controls were used. The far-pair group provided a distance-independent baseline that absorbs any constant common-mode signal. A jitter null was constructed by displacing each source spike by a random offset between 20 and 60 ms, repeated over fifty independent draws.

### Spatial model on real data

To connect the real data to the model, we fit the signed deflection against distance with the same Gaussian kernel form that defines the eLIF field, against a no-coupling (flat) null:

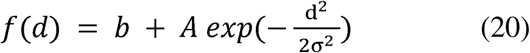

where d is the inter-soma distance, b is a constant offset, A is the amplitude, and σ is the fitted length scale. The fit was performed on distance-binned deflections weighted by their standard errors. We used Bayesian information criterion (BIC, Schwarz, 1978), to compare the null effect and the Gaussian kernel.

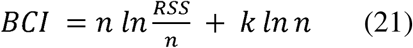

where n is the number of data points, RSS the residual sum of squares, and k the number of free parameters used in the model (**Figure 5g**).

### Single-cell models of the deflection: LIF and eLIF

We asked whether the model used in the rest of the paper reproduces the time course of the measured deflection (**Figure 5h**). The standard LIF model has no field term and predicts no deflection (a flat trace). The eLIF model uses the same coupling as the rest of the paper, the ephaptic field Φ *= α (V_source_ - V_rest_),* which has two effects on a recorded neighbor. First, the extracellular field appears in the whole-cell recording essentially instantaneously, because the pipette reads the extracellular potential directly and volume conduction carries no membrane time constant. Second, the same field polarizes the neighbor membrane, a response that is low-pass filtered by the membrane time constant. We therefore wrote the eLIF prediction for the recorded neighbor STA as the sum of an instantaneous field component and a τ_m_-filtered membrane-polarization component and fit the two coefficients jointly to the data by least squares over ±8 ms. Source spikes were generated by the paper LIF neuron driven just above threshold so that each cell fired sparsely and every averaging window held a single isolated spike; since the LIF resets instantaneously and has no spike waveform, the field was generated by a stereotyped action potential whose width and after-hyperpolarization-to-peak ratio match the measured source action potential (**Figure 5c**), set independently of the neighbor data being fit. The combined eLIF model reproduced the biphasic deflection (coefficient of determination R-squared = 0.69): the instantaneous field accounts for the fast peak and the τ_m_ component for the slower undershoot.

### Statistics for the real data

We applied four tests to the real-data measurements, and their p-values were corrected together by the Holm-Bonferroni procedure: the close-versus-far comparison of the signed deflection (two-sided Mann-Whitney U test), the comparison of the close deflection against the jitter-null (Wilcoxon signed-rank test), the amplitude-against-distance trend (Spearman rank correlation), and the connected-versus-unconnected comparison of the absolute deflection (two-sided Mann-Whitney U test). The agreement between the eLIF field model and the measured deflection was quantified by the coefficient of determination and the Pearson correlation between the model prediction and the data over the ±8 ms window.

## Acknowledgments

This work was supported by a C.V. Starr Fellowship from Princeton University (KBB), This work was supported by a C.V. Starr Fellowship from Princeton University (KBB), and NEI (2R01EY017699; SK), NIH (1R01MH137624; SK), NIMH (1P50MH132642; SK). The funders had no role in study design, analysis, and the decision to publish, or the preparation of this manuscript.

## Author Contributions

K.B.B developed and designed the methodology, modeling, software, analysis. K.B.B and S.K. wrote the paper.

## Data availability

The multi-patch electrophysiology data analysed in this study are part of the Allen Institute for Brain Science Synaptic Physiology dataset (release 2.1), publicly available from the Allen Institute.

## Code availability

The code used to generate the simulations and analyses in this study is available from https://github.com/banaiek/ephaptic_coupling.

---

**Figure S1.**
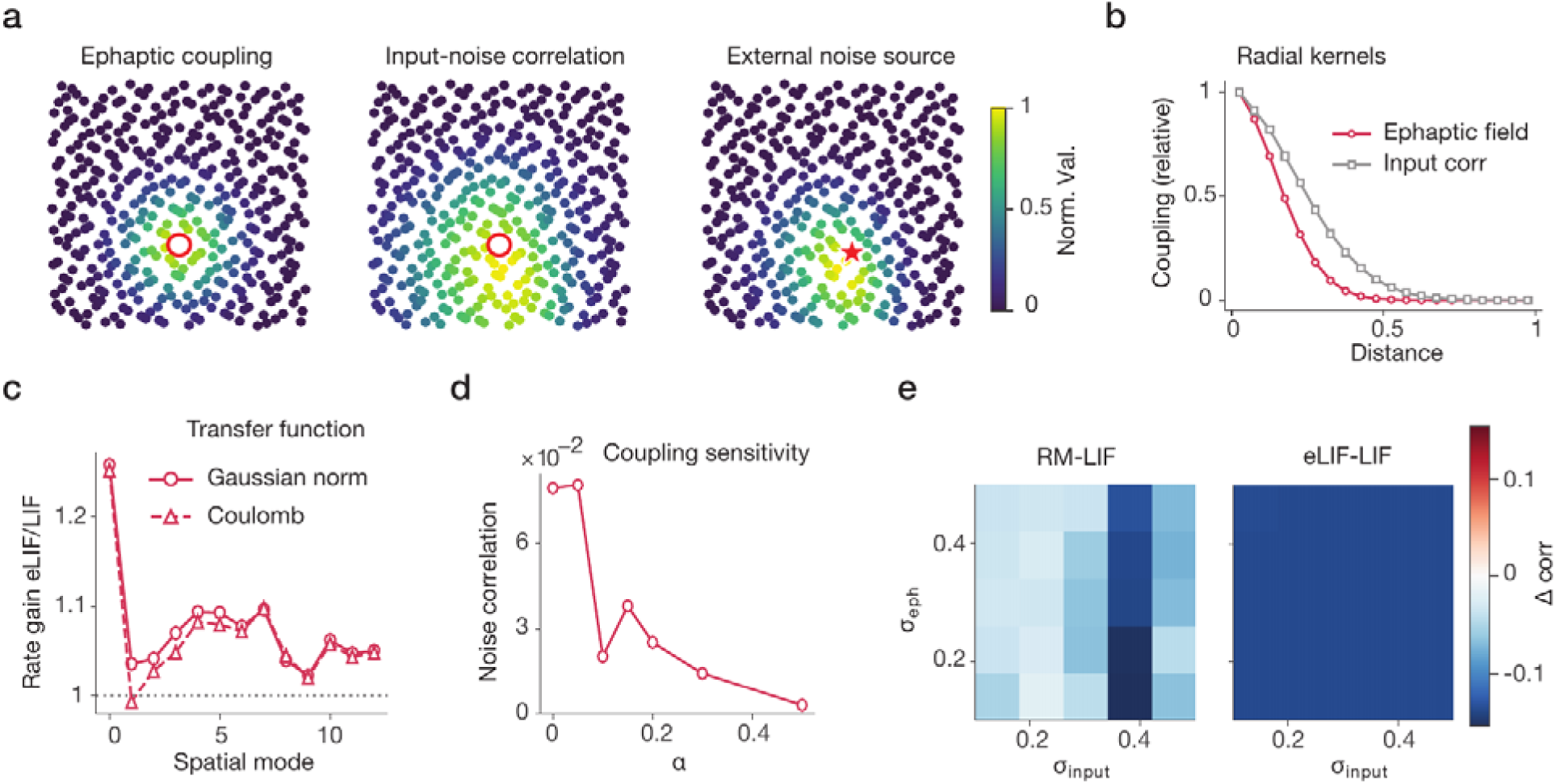
Field characterization of the voltage decorrelation. **(a)** Single-cell spatial footprints for one example neuron (red circle) showing the ephaptic coupling it receives, the correlation of its shared input, and an external noise source. **(b)** Radial coupling kernels averaged over neurons for the ephaptic field and the shared-input correlation. **(c)** The transfer function (the subthreshold voltage gain of the field-coupled relative to the field-free network) across spatial modes, for two different kernel shapes, a normalized Gaussian and a Coulomb kernel. **(d)** Mean pairwise noise correlation as a function of coupling strength α. **(e)** The difference in voltage correlation relative to LIF across a grid of field length scale (σ_eph_) and input length scale (σ_input_). Error bars and shaded bands, SEM.

**Figure S2.**
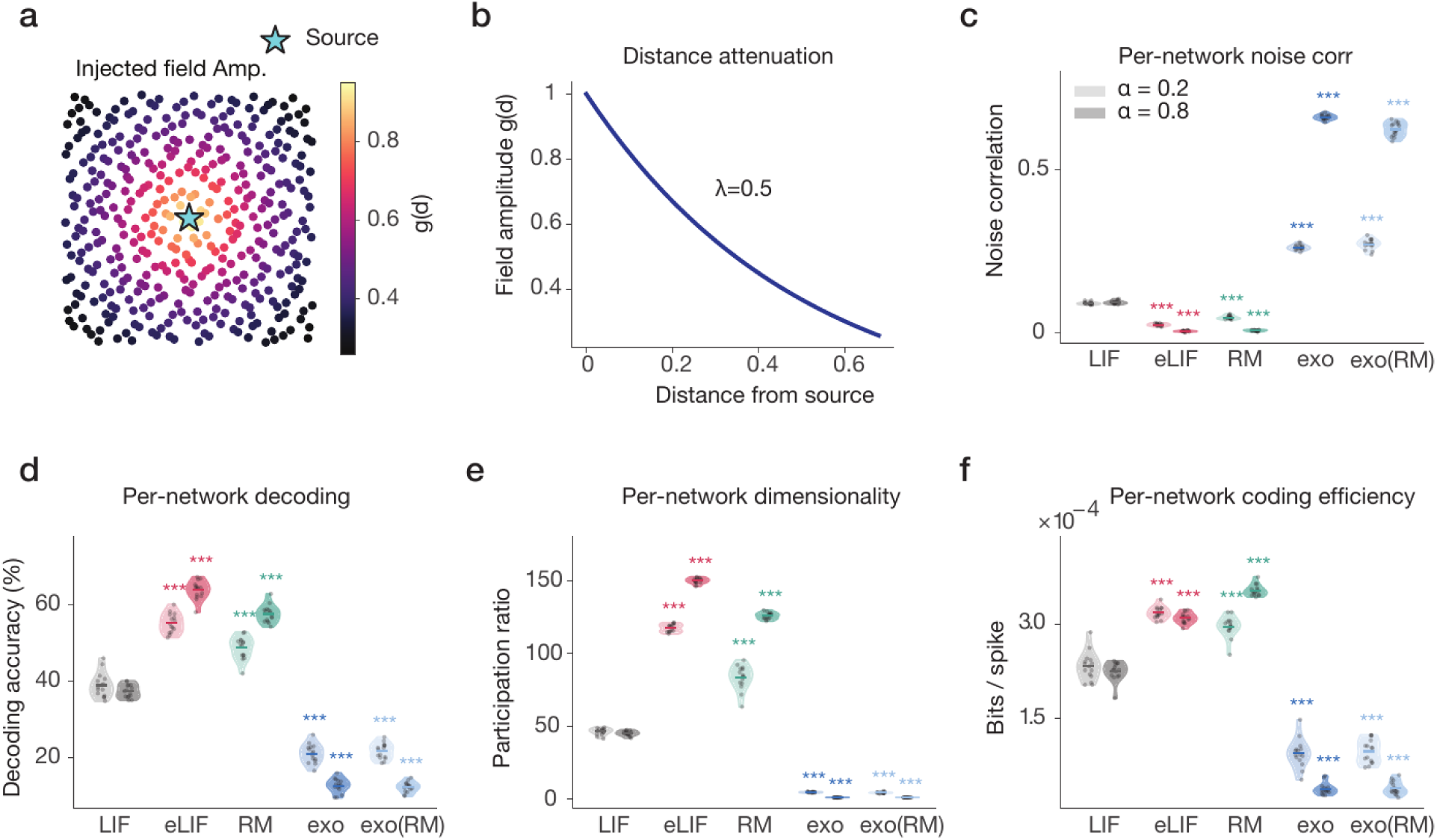
Exogenous field geometry and per-network coding measures. **(a)** Spatial map of the injected exogenous field amplitude across the network relative to the source (star). **(b)** The distance attenuation of the exogenous field. **(c-f)** Per-network measures at weak (α = 0.2, light) and strong (α = 0.8, dark) coupling for **(c)** mean noise correlation, **(d)** decoding accuracy, **(e)** effective dimensionality (participation ratio), **(f)** and coding efficiency (bits per spike) for the five conditions. Each point is one network, and the conditions are LIF, eLIF, RM, exo (exogenous) and exo(RM) (rate-matched exogenous). Dots show individual network seeds. Asterisks show significance level: *p<0.05, **p<0.01, ***p<0.001.

**Figure S3.**
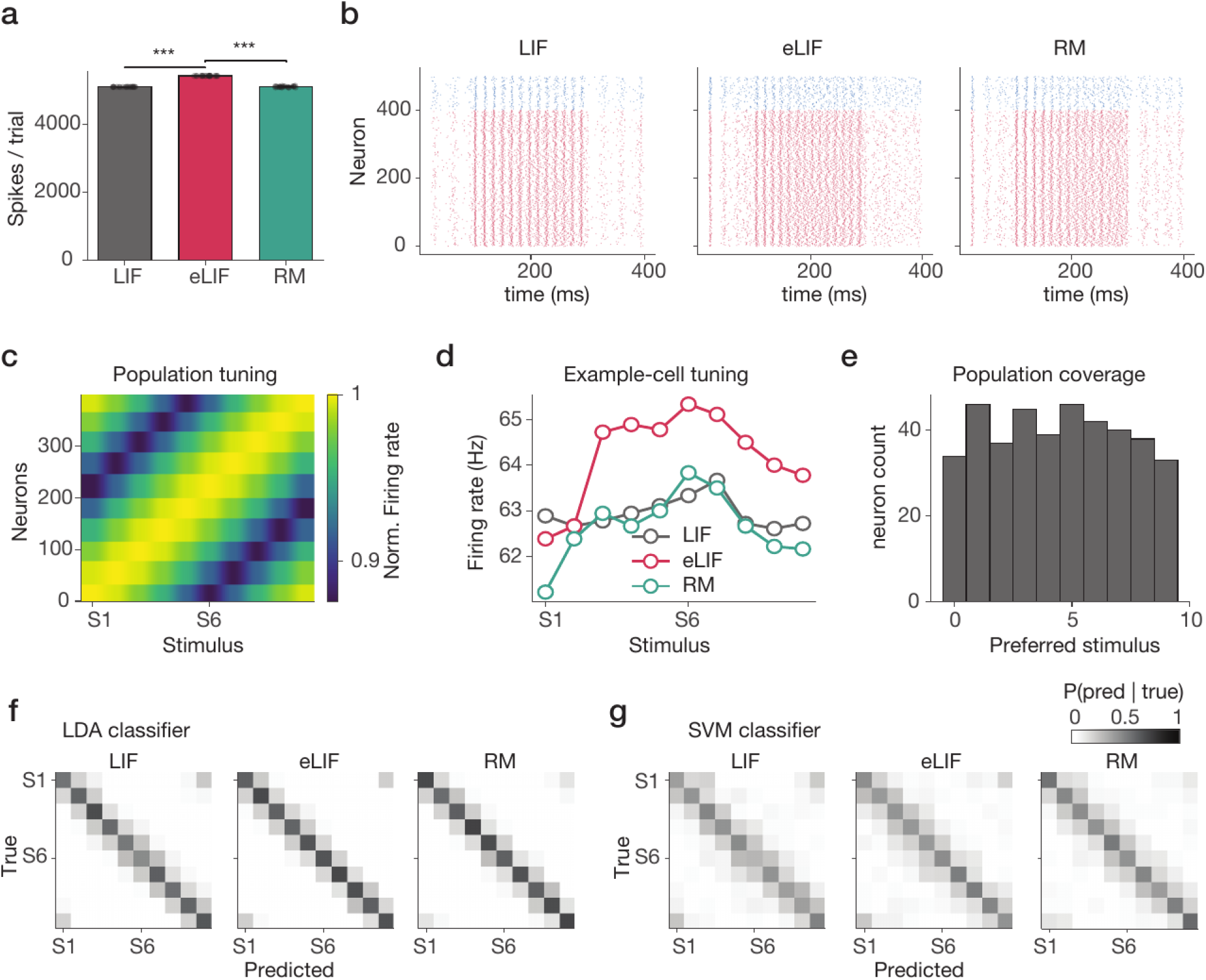
Rate, tuning and decoder controls for the stimulus-decoding task. **(a)** Total spikes per trial for LIF, eLIF and RM. **(b)** Spike rasters for one example trial in each network **(c)** The population tuning map. **(d)** Example single-cell tuning curves for the three networks. **(e)** The population coverage of stimulus space showing the number of neurons preferring each stimulus. **(f,g)** confusion matrices of **(f)** the linear discriminant (LDA) decoder and **(g)** the support-vector machine (SVM) decoder.

**Figure S4.**
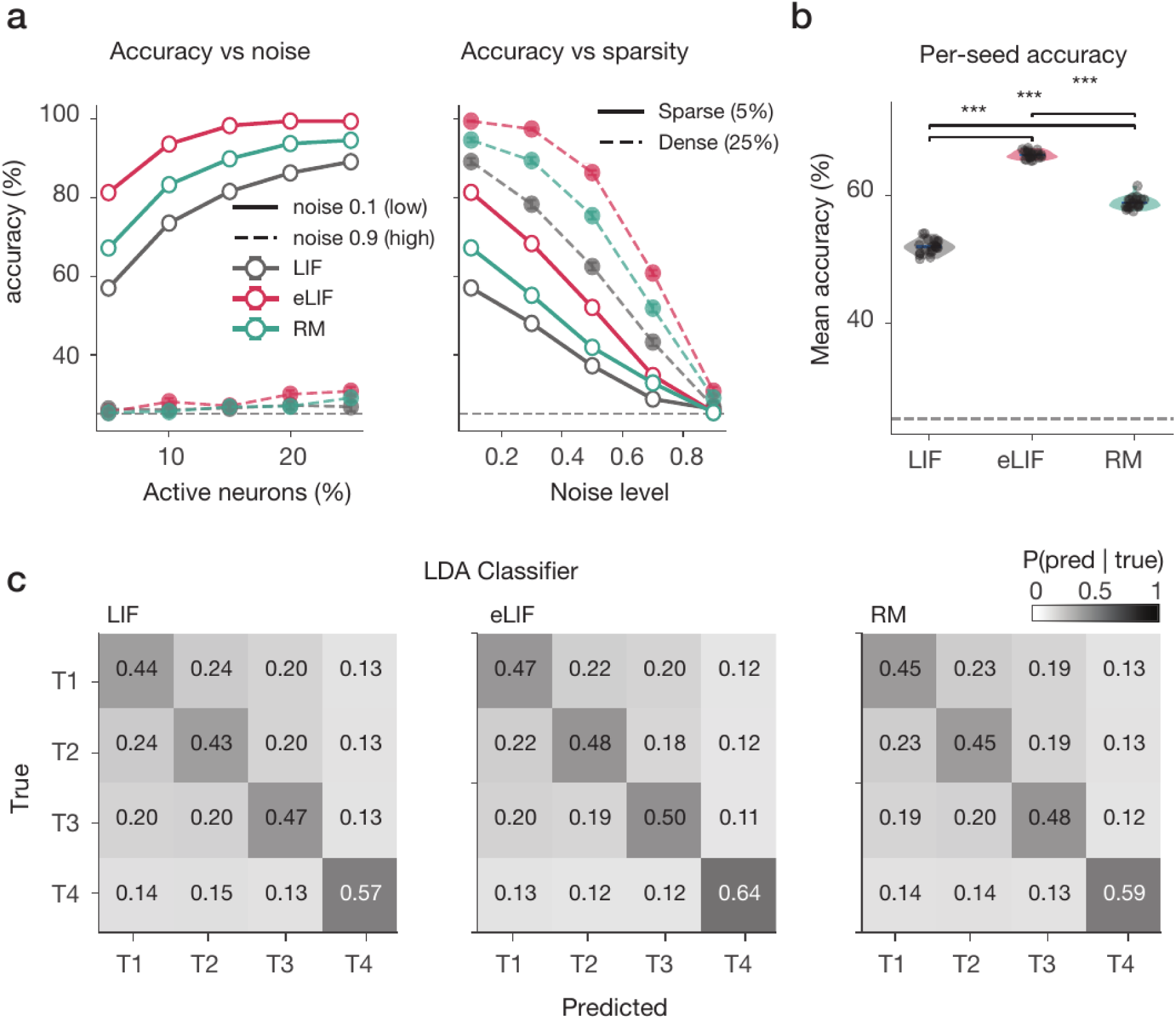
Additional controls for the sparse temporal-coding task. **(a)** Classification accuracy versus the sparsity (left, at low and high noise) and the noise level (right, for sparse and dense activation), for the three networks. **(b)** Per-seed mean accuracy for the three networks. **(c)** Same as in Figure 4g showing Confusion matrices of the linear discriminant (LDA) decoder. Abbreviations: LDA, linear discriminant analysis; T1 to T4, the four temporal patterns; sparse, 5% of neurons active; dense, 25% active; LIF, eLIF and RM as defined in Figure 1. Dots show individual network seeds. Asterisks show significance level: *p<0.05, **p<0.01, ***p<0.001.

**Figure S5.**
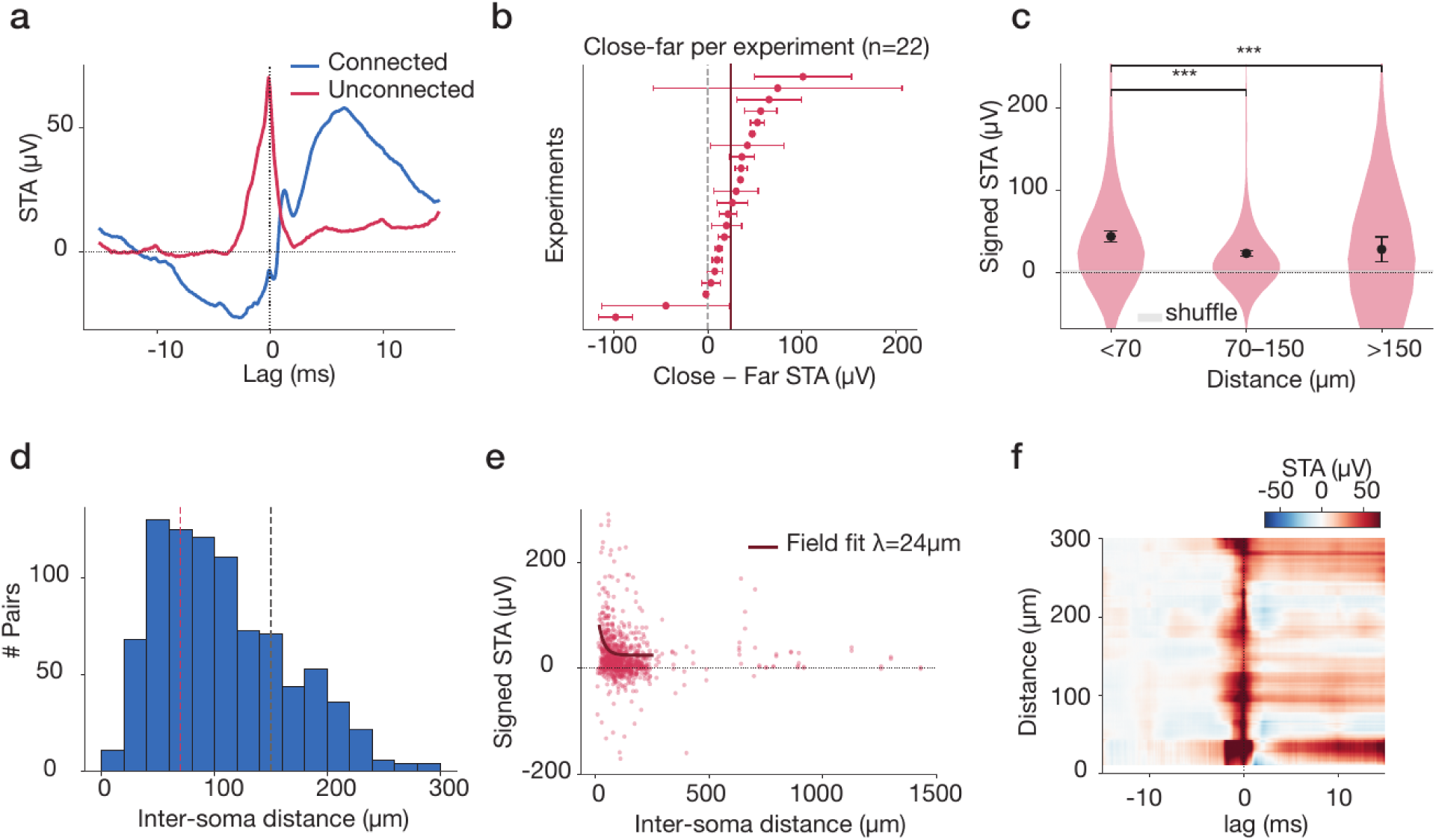
Supporting controls for the cortical spike-triggered signature. **(a)** The spike-triggered average (STA) versus lag for connected (blue) and unconnected (red) pairs. **(b)** The close-minus-far STA difference for each of the 22 experiments. **(c)** The signed STA amplitude by distance bin (less than 70, 70 to 150, and more than 150 µm), with the jitter-shuffle control. **(d)** The distribution of inter-soma distances across pairs. **(e)** The signed STA amplitude as a function of inter-soma distance for all pairs, with the field-kernel fit. **(f)** The STA as a function of both lag and inter-soma. Error barrs indicate SEM across pairs or experiments. Asterisks show significance level: *p<0.05, **p<0.01, ***p<0.001.

